# Identification and Prioritisation of Variants in the Short Open-Reading Frame Regions of the Human Genome

**DOI:** 10.1101/133645

**Authors:** Felix Jackson, Matthew Wayland, Sudhakaran Prabakaran

## Abstract

As whole-genome sequencing technologies improve and accurate maps of the entire genome are assembled, short open-reading frames (sORFs) are garnering interest as functionally important regions that were previously overlooked. However, there is a paucity of tools available to investigate variants in sORF regions of the genome. Here we investigate the performance of commonly used tools for variant calling and variant prioritisation in these regions, and present a framework for optimising these processes. First, the performance of four widely used germline variant calling algorithms is systematically compared. Haplotype Caller is found to perform best across the whole genome, but FreeBayes is shown to produce the most accurate variant set in sORF regions. An accurate set of variants is found by taking the intersection of called variants. The potential deleteriousness of each variant is then predicted using a pathogenicity scoring algorithm developed here, called sORF-c. This algorithm uses supervised machine-learning to predict the pathogenicity of each variant, based on a holistic range of functional, conservation-based and region-based scores defined for each variant. By training on a dataset of over 130,000 variants, sORF-c outperforms other comparable pathogenicity scoring algorithms on a test set of variants in sORF regions of the human genome.

**List of Abbreviations:**

AUPRC: Area under the precision-recall curve
BED: Browser Extensible Data
CADD: Combined annotation-dependent depletion
DANN: Deleterious annotation of genetic variants using neural networks
EPO: Enredo, Pecan, Ortheus pipeline
GATK: Genome analysis toolkit
GIAB: Genome in a bottle
HGMD: Human gene mutation database
Indels: Insertions and deletions
MS: Mass spectrometry
ORF: Open reading frame
RF: Random Forests
ROC: Receiver Operating Characteristics
SEP: sORF encoded peptide
sklearn: Scikit-learn package
SNVs: Single nucleotide variants
sORF: Short open-reading frame
TF: Transcription factor
TSS: Transcription start site
VCF: Variant Call Format file

## Introduction

Comprehensive data on the cellular proteome is required to fully understand a cell’s state. However, the human proteome differs significantly between cell types and even between cells that possess identical genomes. A major contribution to this tissue and cell-type specific variability comes from transcription and translation of regions of the genome previously classified as ‘non-coding’. Most notably, a number of short open-reading frames (sORFs) and protein-like products from the non-coding regions have recently been identified to be translated in humans and other model organisms (Prabakaran *et al*, 2014). sORFs are short lengths of in-frame codons bordered by a start and stop codon, typically between 2-100 codons in length. Although these sORFs are potentially translatable, they have been somewhat disregarded as non-functional gene regions. This view has been challenged in the last decade however, first with the identification of sORFs that were transcribed and conserved between species (Kastenmayer *et al*, 2006; Hanada *et al*, 2007), and then more recently by identifying that many were translated into sORF-encoded peptides (SEPs) (Vanderperre *et al*, 2013; Slavoff *et al*, 2013). Although the number of annotated SEPs remains fairly low, SEPs have already been linked to a range of cellular functions: from metazoan morphogenesis (Hashimoto *et al*, 2008) to mammalian DNA end-joining (Slavoff *et al*, 2014), SEPs are emerging as biologically relevant molecules that should be considered part of the proteome.

sORFs that encode translated peptides had previously evaded detection primarily due to their small size. *Ab initio* gene predictors rely upon sequence specific features such as the Kozak sequence or intrinsic hexamer bias to separate coding from non-coding regions of the genome (Sleator, 2010), and sORFs are generally too small to be enriched in these features. Furthermore, gene prediction algorithms are generally not trained on sORF regions, and many automatically disregard candidate ORFs that are smaller than 100 codons: a cut-off point shown to be arbitrary (Frith *et al*, 2006), but nonetheless widely used. Mass spectrometry (MS) based methods have failed to detect SEPs because of their low abundance as it is technically challenging to detect peptides of lower abundance and size.

However, as sORFS are becoming recognised as functionally important determinants of cell state, these technical challenges are being overcome. This is exemplified by tools such as sORF finder (Hanada *et al*, 2009), a freely available online tool that can predict the locations of sORFs with high-coding potential. The advent of ribosome profiling (Ingolia *et al*, 2009) represents an important advance, as it provides evidence of mRNA transcripts in complexes with ribosomes in a translation initiation state. Although additional regulatory steps prevent all mRNA-ribosome complexes being translated to functional peptides, it is at least a useful indicator for translation. A comprehensive list of 58,137 non-overlapping sORFs identified by ribosome profiling of a human carcinoma cell line has been made freely available online at sORFs.org (Olexiouk *et al*, 2016), and other similar data is emerging elsewhere.

Just as technical innovations will likely reveal the prevalence of SEPs in the human proteome, mutations in sORF regions of the human genome will emerge as important determinants of disease. Indeed, there is already evidence that a set of mutations in the upstream sORF of the hairless homolog gene (HR) cause translation of a SEP that is associated with Marie Unna hereditary hypotrichosis (Wen *et al*, 2009). To identify potentially pathogenic mutations in a high-throughput manner, two tools are required. First, a variant caller is needed to identify all sites in the human genome at which mutations have occurred. However, it is estimated that variants occur for one in every thousand nucleotides in *Homo sapiens*, and the vast majority of these mutations are benign. The second tool required to link identified variants to disease is therefore a pathogenicity prediction tool, which prioritises variants in terms of their likelihood of being pathogenic.

Variant calling is the process of separating signal from noise in sequencing experiments: a 50x paired-end whole genome sequence produces roughly 1 billion reads, which contain on average 3 million variants. This requires algorithms that can accurately and reliably distinguish true variants from experimental artifacts and mapping effects. Instrumental to this process are various quality and context-dependent metrics, which are assigned to each base during sequencing and subsequent processing. The variant caller must then evaluate the likelihood that the observed variant represents the true genotype, based on the distribution of these metrics, using ‘truth’ data such as dbSNP (Sherry *et al*, 2001) as a reference. Importantly, these pattern recognition techniques must work on diverse regions of the genome, where significantly different distributions of each metric are observed. For example, variants within repeat regions have proved particularly challenging to validate (Treangen and Salzberg, 2011), and bases in these regions will have distinct base quality score distributions to those in exons. Although germline variant callers are now highly accurate over most genomic regions, more work is needed to improve their performance in non-coding regions, which are enriched in repetitive elements. Furthermore, excluding exons there has been little effort to systematically compare the performance of these algorithms on specific genomic features. For example, sORF regions present a particular challenge to variant callers, as a significant number lie within poorly annotated, highly repetitive regions of the genome.

Having called variants in sORF regions, their functional effect can be predicted. Numerous tools to predict the pathogenicity of variants have already been devised. Historically, these tools have focused on exonic regions, and have used exon-specific features such as the folding of the predicted protein to predict pathogenicity (Choi and Chan, 2015). Due to the lack of functional information, these tools perform particularly poorly on non-coding regions of the genome. More recently, some algorithms have been adapted for use on the whole genome, including non-coding regions, and use more general variant annotations such as inter-species conservation scores and local chromatin states. The best performing of these use supervised machine-learning algorithms (Kircher *et al*, 2014; Quang *et al*, 2015), where the classifier learns to make accurate predictions from training data. The performance of these algorithms depends directly on the training set used, and so far pathogenicity predictors have only been trained on exon-specific or whole genome data, which generally worsens their feature-specific performance compared to specialist predictors. No pathogenicity predictors have yet been optimised for variants in sORF regions.

Here we present the first steps to be taken towards improving the accuracy and reliability of variant callers and variant prioritisation algorithms for sORFs. The performance of variant calling pipelines in sORF regions was investigated, and an optimised set of variant calls in these regions was collected by taking the intersection of variant sets from each caller. Secondly, a classifier was built that can predict the pathogenicity of known variants in sORF regions with high accuracy. The classifier uses Random Forests, a supervised machinelearning algorithm that is well suited to predicting outcomes which lack reliable predictor variables (Breiman, 2001). It is likely that a number of obscure factors interact to determine the pathogenicity of a genomic mutation; when there are no obvious rules to predict an outcome, Random Forests instead combines a large number of ‘weak’ rules, and achieves accuracy by a strength-in-numbers approach. By training on a holistic range of sORF specific features, and using a machine-learning algorithm that is known to perform well on datasets with large feature sets, a pathogenicity classifier is produced that has better precision and sensitivity than other whole-genome pathogenicity predicting tools.

## Results

### Calling variants in the NA12878 whole genome

Before investigating pathogenicity scoring algorithms in sORF regions of the genome, an accurate list of mutations in these regions is required. The performance of commonly used germline variant calling pipelines was therefore evaluated, by comparing the resulting Variant Call Format files (VCF) to a VCF file containing high-confidence variant calls. In 2014, the Genome in a Bottle consortium (GIAB) published a set of high-confidence variant calls for the NA12878 individual (Zook *et al*, 2014) which was obtained by integrating datasets from a variety of read-mapping and variant calling algorithms, then manually arbitrating discordant data. This VCF is widely used to benchmark variant calling pipelines, as it is currently the only ‘gold-standard’ set of variant calls freely available.

The NA12878 genome reads were obtained in FastQ format, from an Illumina paired-end sequencing experiment at 50x depth. Variant calling pipelines were first evaluated on the whole genome, excluding the highly repetitive regions for which Zook *et al* (2014) could not produce high-confidence calls (approximately 13.4% of the genome). Through aligning, processing, then variant calling with four different variant callers, four VCFs were produced which could be benchmarked against the high-confidence GIAB VCF. The sequence of steps that were performed is illustrated in Figure 1.

**Figure 1:**
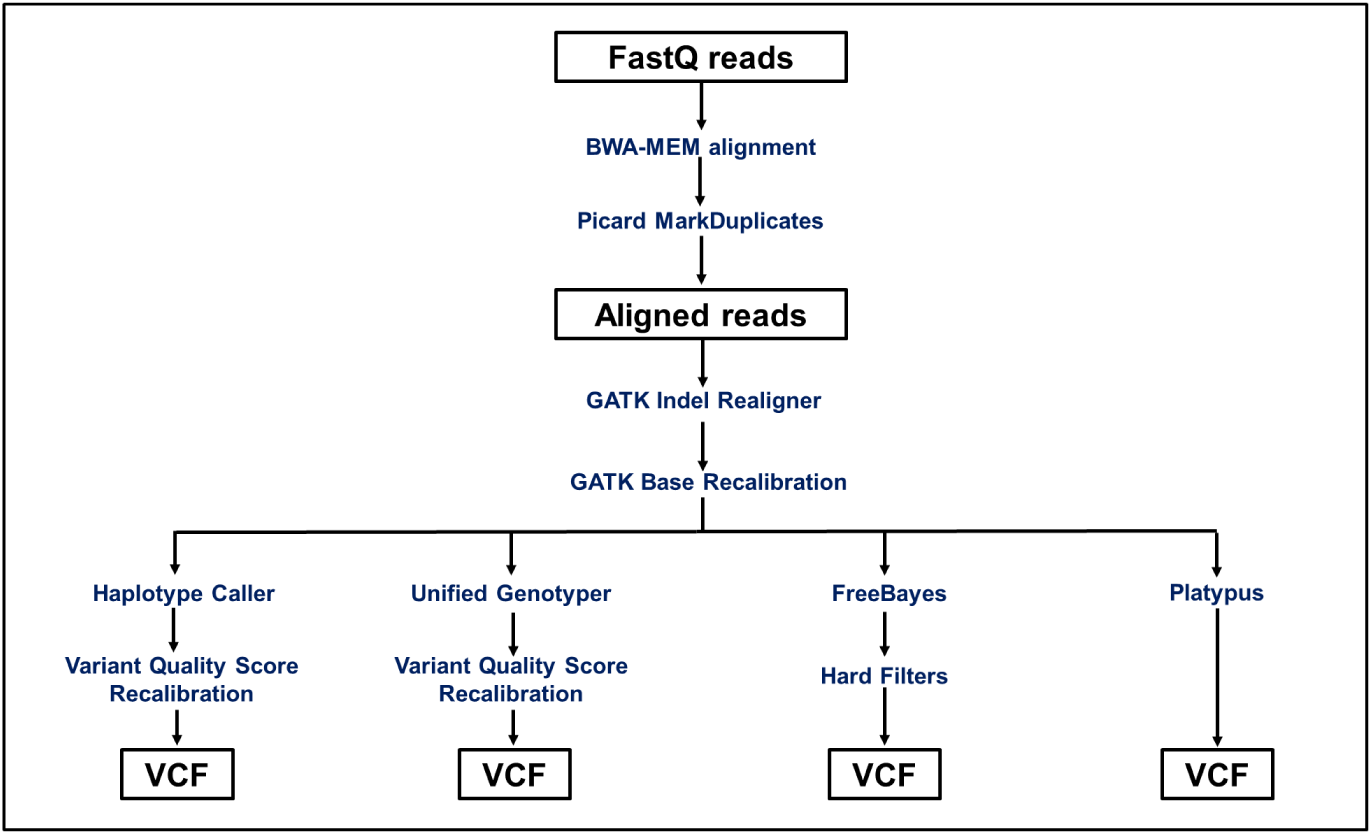
The variant calling pipeline used to call variants in the NA12878 genome. The same aligned, base-recalibrated reads file was tested on all four variant calling algorithms, to produce four distinct lists of variants in the NA12878 genome. The individual steps taken are described in detail in Materials and Methods.

The resulting VCFs from each variant caller were compared to the gold standard GIAB VCF using Real Time Genomics’ vcfeval tool (Cleary *et al*, 2015). This tool provides a more sophisticated way to compare the calls in two VCF files than previously existed. The naive way of simply checking if the same variant type is present at the same genomic location in each VCF does not account for the problem that there are multiple ways of representing the same variant; one multinucleotide polymorphism in the first VCF may be called as several single nucleotide polymorphisms in the second. Briefly, the vcfeval algorithm takes the call from each VCF at a given genomic coordinate, and reconstructs the transition to the reference genome. The reconstruction that maximises true positives and minimises false positives and false negatives is the one chosen. This global optimisation is now accepted as the optimal means of comparing variants, despite the disadvantage that indels and SNVs cannot be considered separately.

The standard method of comparing called variants to a truth dataset is to generate Receiver Operating Characteristic (ROC) curves. However, on imbalanced datasets ROC curves are less informative than precision-recall curves (Saito and Rehmsmeier, 2015). Precision-recall curves show how precision varies with sensitivity (also known as recall), where precision and sensitivity/recall are defined as:

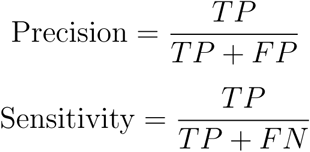

where *T P* is the number of true positives, *F P* the number of false positives, and *F N* the number of false negatives. The harmonic mean of precision and sensitivity, *F*_1_, gives a useful measure of overall accuracy, and is defined as:

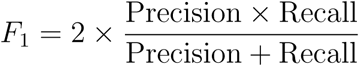

The curves are obtained by iteratively increasing the threshold at which data is classified into one of two bins, then calculating the precision and sensitivity at each step. To create precision-recall curves for sets of variant calls, the threshold used was the Phred-scaled quality score of each variant, which is generated for each call by the variant caller; this threshold was iteratively increased over the entire range of values in the dataset, and at each iteration all variants with a quality score above the threshold were called, while all those with a quality score below were discarded.

The optimal means of evaluating variant calls in a VCF file is to plot these precision-recall curves, and compare the area under the precision-recall curve (AUPRC) between variant callers, as suggested by Hwang *et al* (Hwang *et al*, 2015). The maximum F-measure (*F*_1_ score) can be interpreted as the maximum weighted average of precision and sensitivity, and was therefore used here as an additional metric to assess classifier performance. Precision-recall curves for each variant caller are presented in Figure 2, and the corresponding AUPRC and the maximum F-measure are shown in Table 1.

**Table 1:**
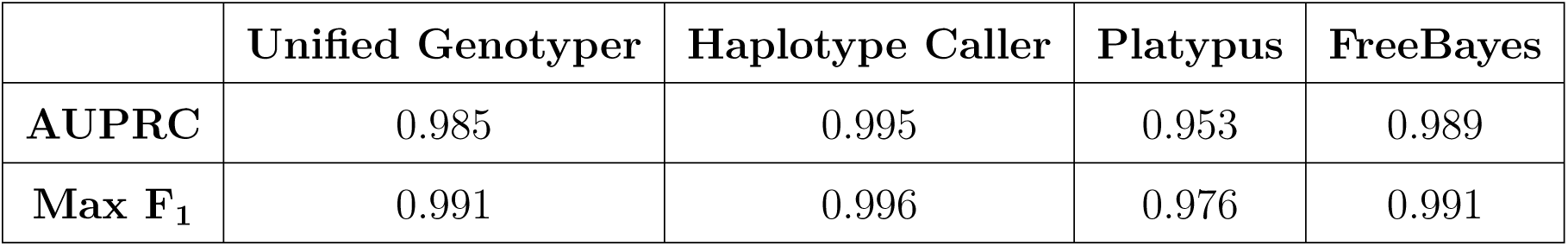
The area under the precision-recall curve and the maximum F-measure for each variant caller.

|  | Unified Genotyper | Haplotype Caller | Platypus | FreeBayes |
| --- | --- | --- | --- | --- |
| **AUPRC** | 0.985 | 0.995 | 0.953 | 0.989 |
| **Max $F_1$** | 0.991 | 0.996 | 0.976 | 0.991 |

**Table 2:** A comparison of AUPRC and F-measure for each set of variants in sORF regions.

|  | Unified Genotyper | Haplotype Caller | Platypus | FreeBayes |
| --- | --- | --- | --- | --- |
| **AUPRC** | 0.702 | 0.714 | 0.720 | 0.739 |
| **Max $F_1$** | 0.817 | 0.828 | 0.836 | 0.837 |

**Figure 2:**
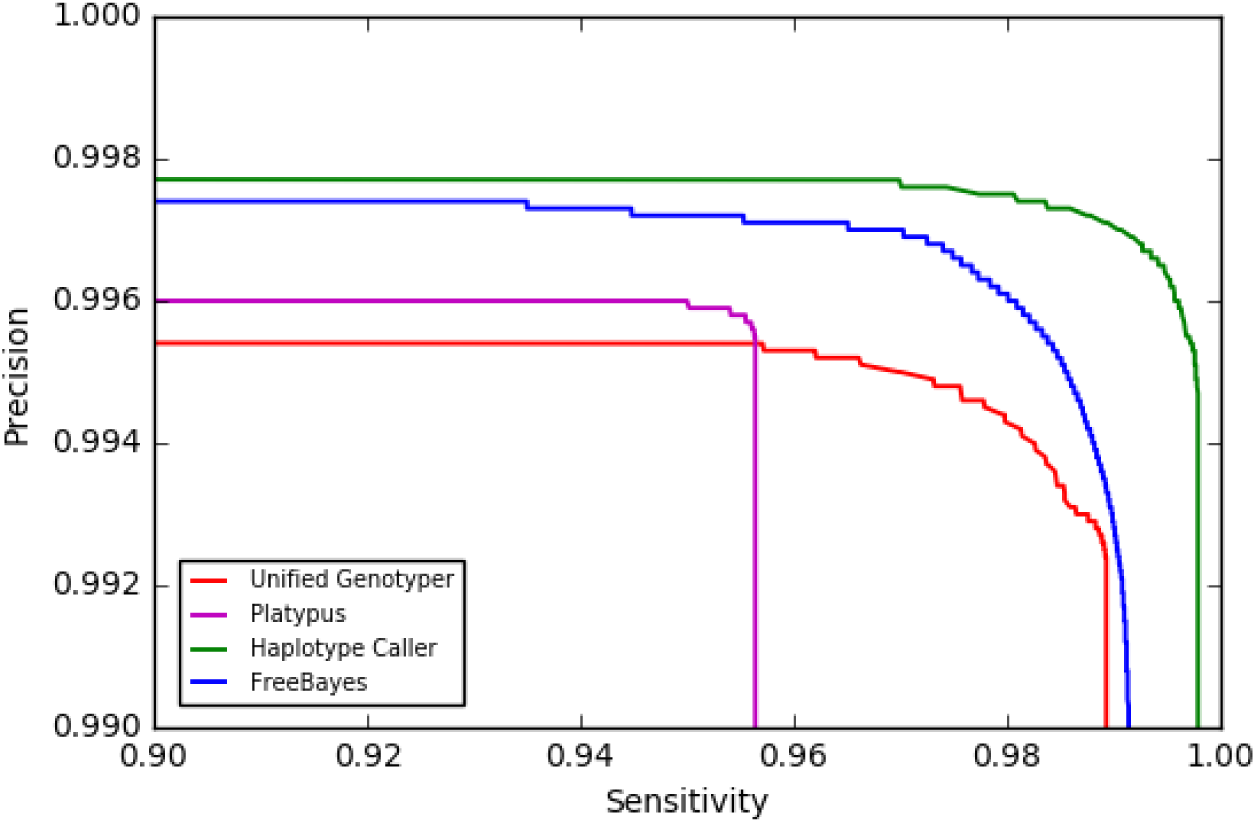
Precision-recall curves facilitate comparison of each variant calling algorithm over the complete range of precision and sensitivity values. The precision-recall curves for the resultant VCFs of each variant calling algorithm were generated using vcfeval, which compared each to the GIAB gold standard VCF. To make the differences in each variant caller’s precision and sensitivity values easily visible, their values over only a small subset of the graph are shown. This was the only region of the graph that differed between variant callers.

These four variant callers demonstrate similarly high precision and sensitivity, and comparable AUPRC and F-measures. The small variations seen are relevant nonetheless; considering approximately four million variants are called in each VCF, the difference of one percentage point could mean losing up to 40,000 variants. Haplotype Caller demonstrates the highest AUPRC and F-measure, and the two scores correlate well as performance metrics.

### Calling variants in short open-reading frames

In order to investigate tool performance in sORF regions, a comprehensive list of human sORFs was needed. The full set of sORFs identified by ribosome profiling of human carcinoma cell line HCT116 was therefore downloaded from sORFs.org (Olexiouk *et al*, 2016). In total, 58,137 non-overlapping sORFs were downloaded, and their relative distribution across different regions of the genome can be seen in Figure 3.

**Figure 3:**
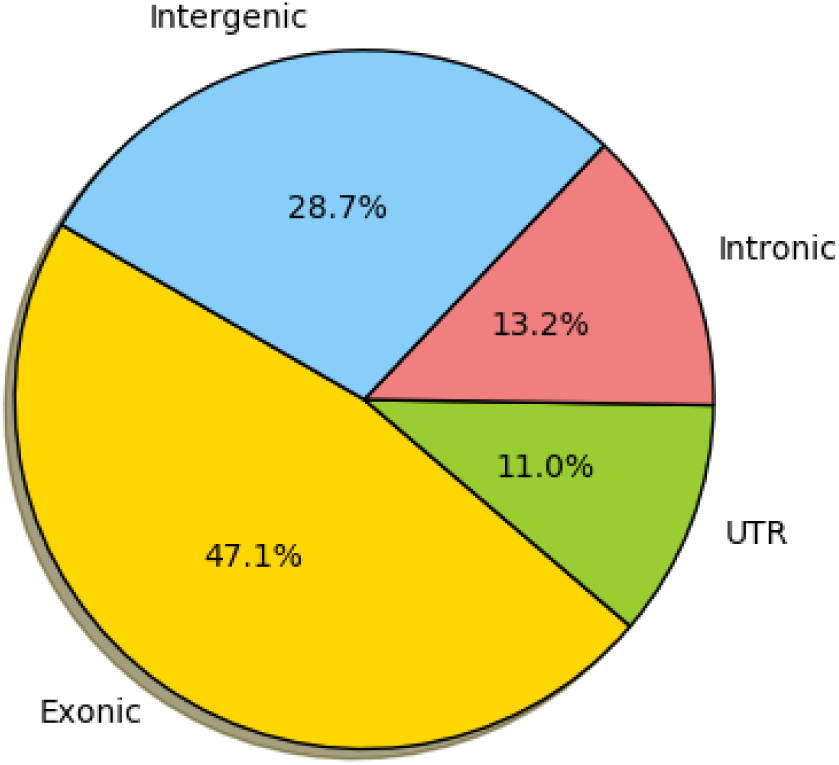
The fraction of sORFs in each genomic region. Exonic regions are defined as regions between a transcription start and stop site which are known to be translated. The UTR regions are counted together here, and are defined as the untranslated gene regions upstream of the first exon or downstream of the last. Intronic regions are the regions bounded by a transcription start and stop site that are spliced out of the mRNA transcript. Intergenic regions consist of all DNA between known coding genes. There are 58,137 non-overlapping sORFs in this dataset.

The performance of each variant caller was then compared within these sORF regions, by specifying the regions of interest over which vcfeval should compare calls to the truth dataset. Precision-recall curves were used again to compare variant caller performance in these sORF regions (Figure 4).

**Figure 4:**
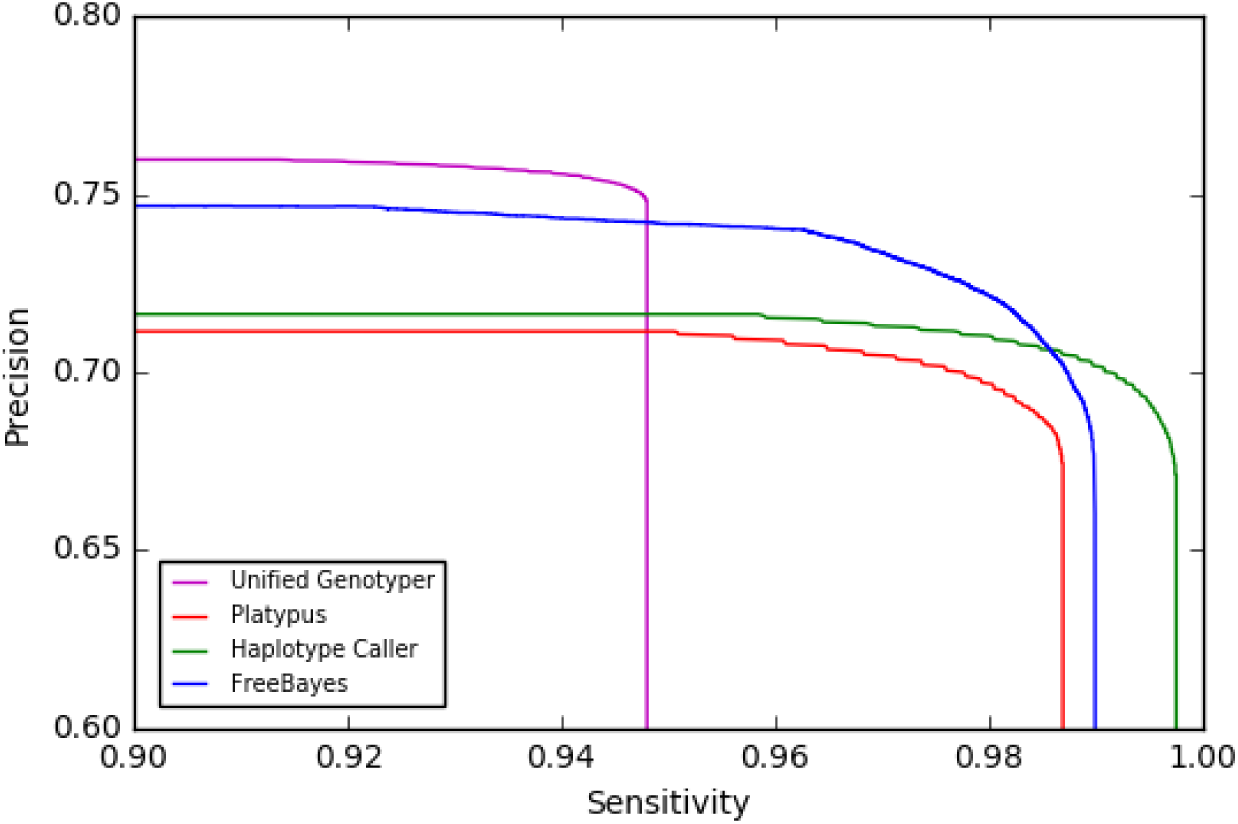
A comparison of each variant caller’s precision and sensitivity values in sORF regions. All variant callers demonstrate greatly reduced precision in variants called, due to the presence of many more false positives (variants that are present in the call set but not in the GIAB truth VCF). Again, only the region of the graph where the variant caller’s precision and recall values differ are shown.

Here, precision and sensitivity is compared over all sORF regions, some of which overlap the repeat regions that were excluded in the previous genome wide comparison. An important contributing factor to the low precision of all VCFs is therefore the poor coverage of repeat regions in the GIAB gold-standard VCF. Indeed, 5.0% of sORF regions used here were not in the GIAB high-confidence call regions. Because the truth dataset lacks full coverage in these regions, many variants called as false positives here may in fact be true variants that are not present in the GIAB ’truth’ variant set. This is a confounding factor when evaluating the performance of variant callers in sORF regions.

It is notable nonetheless that Haplotype Caller performs worse than Platypus and FreeBayes variant callers here; this may be because Haplotype Caller deals less well with the newly-introduced repeat regions. In an effort to improve this precision, the intersection of the four sORF specific variant sets was taken. The resulting variant set contained 197,126 variants, compared to the average of 214,535 in the original variant sets. Removing the 17,409 variants (8.1%) that were only called by a subset of the variant calling algorithms improved the precision of the call set, at the expense of slightly reduced sensitivity: the maximum F-measure increased to 0.842, but the AUPRC remained low at 0.718. By extensive processing and score recalibration pre- and post-variant calling, and then taking the intersection of the variant sets and thus removing 8.1% of the more dubious variants, a set of sORF-specific variants were obtained, whose accuracy is now only limited by the absence of a high-confidence callset for the entire genome.

### Variant prioritisation

Having collected an accurate list of variants, a classifier was developed to prioritise these variants by their predicted deleteriousness. To train the classifier, ‘truth’ datasets (where the pathogenicity of a variant is known, or at least asserted to a sufficient confidence level) were obtained from three distinct sources. Pathogenic mutations were obtained from the Human Gene Mutation Database (HGMD) (Stenson *et al*, 2014), which contains a list of inherited mutations that have been identified as pathogenic in peer-reviewed literature. Only the 25,013 variants from HGMD contained within the sORF regions (as identified previously) were used. The ClinVar database aggregates variants submitted manually from research groups globally, each with a phenotypic interpretation, and a manually curated clinical significance score. The database was filtered to only include variants contained within sORF regions and classified as ‘benign’ or ‘pathogenic’, resulting in 4,985 pathogenic variants, and 1,960 benign variants being taken from ClinVar.

A third dataset of exclusively benign variants was downloaded from (http://krishna.gs.washington.edu/members/mkircher/download/CADD/v1.3/training_data) with the kind permission from Dr. Martin Kircher. This dataset contains benign variants inferred by identifying differences between the current human reference genome (hg19) and the inferred human-chimp ancestral genome, as identified in the Ensembl EPO 6-way Primate alignment. Variations in the current reference genome compared to our ancestral genome can be assumed to be benign because they have not undergone negative selection since humans evolved from the last human-chimp common ancestor. The subset of these variants that are present in sORF regions was taken, from which 100,000 were randomly sampled for the truth dataset.

Each variant in these datasets of known pathogenic and benign mutations was then annotated with a variety of scores using Annovar (Wang *et al*, 2010). This takes variant annotations from online datasets, and applies them in a base-specific manner. These were primarily conservation-based (such as PhastCons and PhyloP), functional-effect predictions (such as GWAVA and GERP++) and region-based scores, such as chromatin modifications or transcription factor binding. The full list of annotations, and the protocol followed within Annovar can be found in Materials and Methods. Importantly, a ‘pathogenic’ mutation here is defined rather loosely as any variant that is causally linked to a disease. The presence of that mutation alone may not be the sole cause of the associated disease, but the presence of the pathogenic mutation will increase the likelihood of the disease occurring. This definition reflects the classification criteria used to submit variants to the HGMD and ClinVar truth datasets, so unfortunately a more precise definition cannot be used.

The Random Forests supervised learning algorithm was used to build a classifier that is optimised for variants in sORF regions. This algorithm learns a set of rules with which to predict a specified outcome variable, by training on a truth dataset for which the outcome variable is known. High predictive accuracy is obtained by pooling the votes of multiple decision trees formed on random subsamples of the data. Random Forests (RF) was used as it is relatively robust to overfitting, and is well suited to ‘small n, large p’ problems, where the number of parameters (p) is relatively high compared to the sample size (n).

The truth dataset was randomly partitioned into training data (75%) and test data (25%), and the classifier used the training data to learn a set of rules that could be used to predict pathogenicity. These rules were then applied to the remaining 25% test data, and each variant was classed as pathogenic or benign. After optimising the tuning parameters (see Table 6 Materials and Methods), this algorithm correctly classified 99.47% of variants held out in the test set.

An RF regression model was also tested, which used identical training data and parameter tunings; the only difference being the outcome was predicted as a continuous value between zero and one, where a higher value represents increased likelihood of the variant being pathogenic. This facilitated the use of precision-recall curves to more accurately compare this model to other pathogenicity scorers, as these other scorers also output continuous scores that represent the likelihood of being pathogenic. The pathogenicity scoring tool developed here (sORF-c) was compared to the pathogenicity predicting algorithms that currently perform best across the whole genome, including in non-coding regions: CADD (Kircher *et al*, 2014), DANN (Quang *et al*, 2015) and FATHMM (Shihab *et al*, 2015). The precision-recall curve, AUPRC and F-measure of sORF-c compared to the other scorers is shown in Figure 5 and Table 3.

**Table 3:**
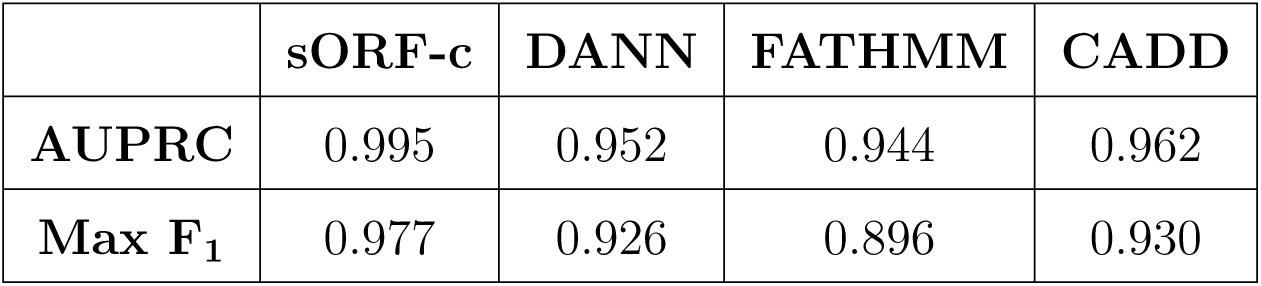
A comparison of AUPRC and F-measure for each pathogenicity scoring algorithm.

**Figure 5:**
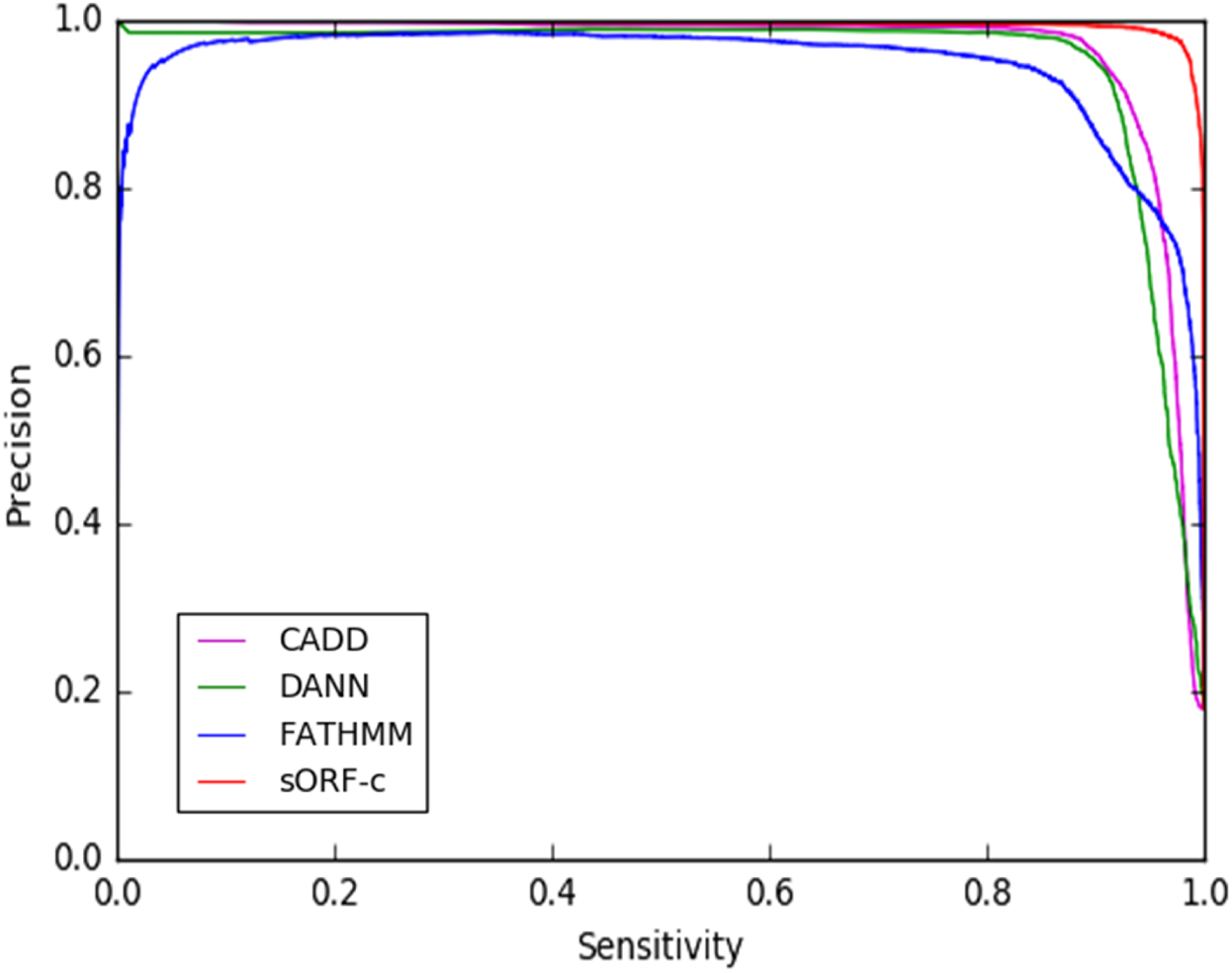
A precision-recall curve comparing the performance of the best currently existing whole-genome pathogenicity scorers to sORF-c. On the test dataset, sORF-c achieves higher precision than other pathogenicity scorers, at higher corresponding sensitivity values. Precision-recall curves were generated for each by iterating through the range of pathogenicity scores output by the scorer, and using each iteration as the threshold above which variants are classed as pathogenic. Precision and sensitivity at each threshold value can thus be calculated.

As demonstrated here, sORF-c outperforms the other available options when calling variants in sORF regions from the test set, demonstrating a higher AUPRC and F-measure. This distinction between the scoring algorithms is clearer when comparing precision with a fixed sensitivity. For example, to obtain a sensitivity of 98%, 98% of all pathogenic variants must be identified. All classifiers can obtain this sensitivity (for example by simply calling all variants pathogenic), but at variable reductions in precision, as more false negatives are introduced. The precision of each classifier at a fixed sensitivity of 98% is compared in Figure 6. Evidently, sORF-c demonstrates greater precision with this constraint, meaning far fewer false positives must be called in order to capture 98% of the true pathogenic variants.

**Figure 6:**
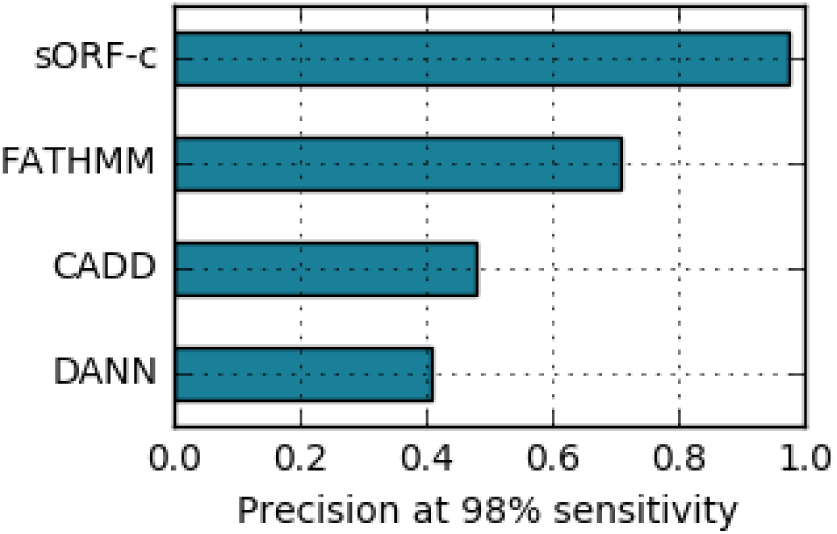
The precision of each scoring algorithm when sensitivity is fixed at exactly 98%. Precision values (as a fraction): DANN = 0.407; CADD = 0.478; FATHMM = 0.708; sORF-c = 0.973

Given this truth dataset, it was also possible to calculate the optimal threshold score above which variants should be classed as pathogenic. This was done for each pathogenicity scoring algorithm, by identifying the threshold that results in the largest F-measure (Table 4).

**Table 4:** The threshold value that results in the highest F-measure for each pathogenicity scoring algorithm compared here.

| Classifier | sORF-c | FATHMM | CADD | DANN |
| --- | --- | --- | --- | --- |
| Threshold | 0.565 | 0.892 | 14.972 | 0.925 |

### Validating the sORF classifier on new data

To investigate the general utility of sORF-c, its performance on new datasets was tested. Unfortunately, this important step was limited by the paucity of manually curated databases with proven pathogenicity classifications. Nevertheless, three further tests were performed.

Firstly, the benign and pathogenic variants from ClinVar were excluded from the training set, and then the trained classifier was used to predict the pathogenicity of each of the excluded variants. This resulted in an error rate of 24% (where error rate is defined as the sum of false positives and false negatives, divided by the total number of calls), suggesting that the ClinVar database provides crucial training data to the model. Secondly, the pathogenicity of the 453 variants within ClinVar classified as ‘probably pathogenic’ and ‘probably benign’ was predicted with the classifier. These variants were classified with a modest error rate of 11%.

In an attempt to identify reasons for the contrasting performance of sORF-c on the training data versus new probable variants in the ClinVar database, the distribution of feature scores in the training datasets was compared. The score distribution of many of the features differed substantially between the two distinct sources of benign variants, whereas they were largely similar for the two pathogenic datasets (data not shown). These underlying differences between benign variants inferred by aligning to the ancestral genome and those reported clinically in ClinVar could be the cause of the sub-optimal predictive performance of sORF-c, when tested on new data from ClinVar. However, it should be noted that these variants were classified as ‘probably’ pathogenic or benign, so this extra degree of uncertainty in their classification may be a confounding factor here.

Finally, the pathogenicity of the intersection of variants called in the NA12878 genome were classified. The classifier predicted 195,090 benign variants, and 2036 pathogenic variants. Although there is no way of validating this prediction, it seems a reasonable estimate; the vast majority of genomic variants are benign, and moreover the NA12878 female has no known diseases. Of the called variants classified as pathogenic, 78 were already in the HGMD database, and 17 were in the ClinVar database classified as pathogenic.

### Variable importance in the Random Forests regression model

In order to understand the contribution that each feature makes in the RF decision model, variable importances were calculated (Figure 7). The Python SciKit-learn package used here (see Materials and Methods) calculates feature importance using the Gini importance score, which is formally defined in (Louppe *et al*, 2013), but can be interpreted as the relative (standardised) importance of each feature in the model.

**Figure 7:**
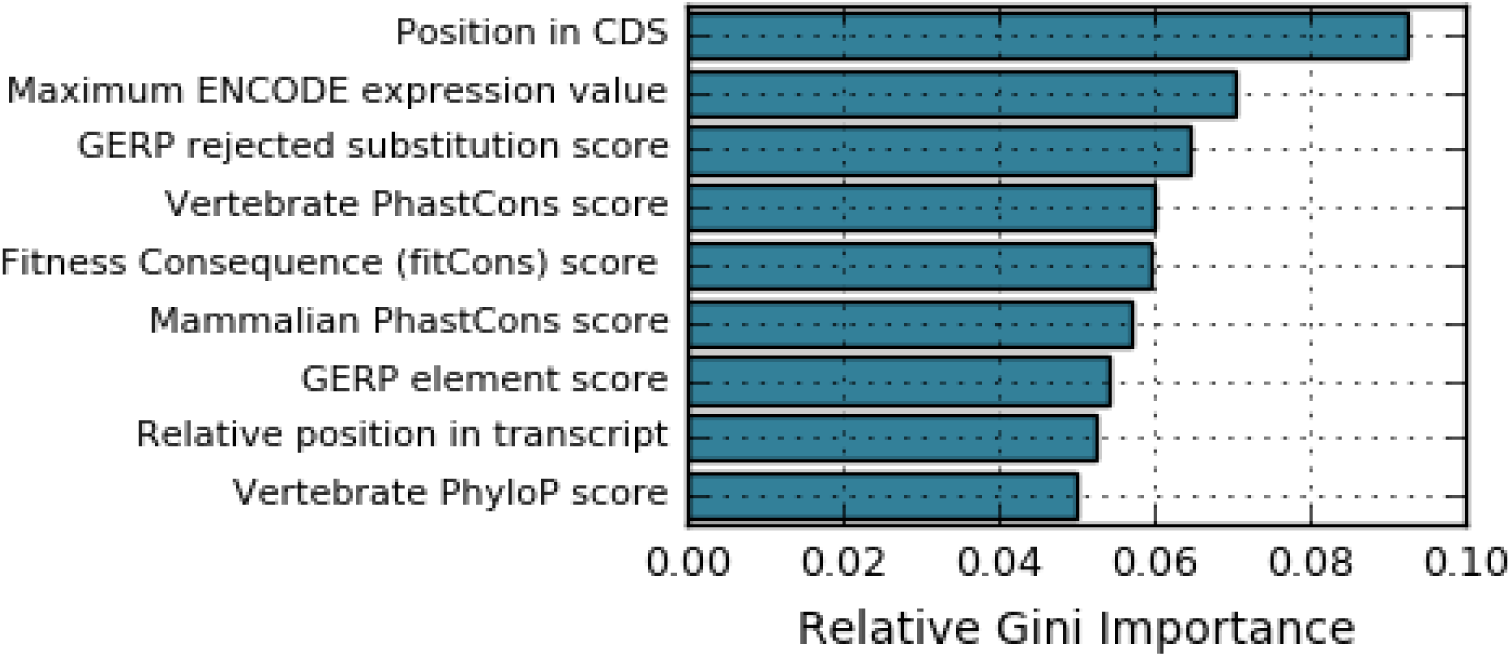
The relative importance of features in the RF regression model. The nine features with the highest importance scores are shown here. All values are standardised by expressing as a ratio of the total Gini importance. Table 5 in Materials and Methods contains more details on each feature.

The gradual decrease in importance of each feature shown in Figure 7 continues with the same trend for the rest of the features in the dataset, demonstrating that no single feature dominates the decision making process. Instead, a holistic range of functional, region-based and conservation-based features are taken into account in this model.

**Table 5:**
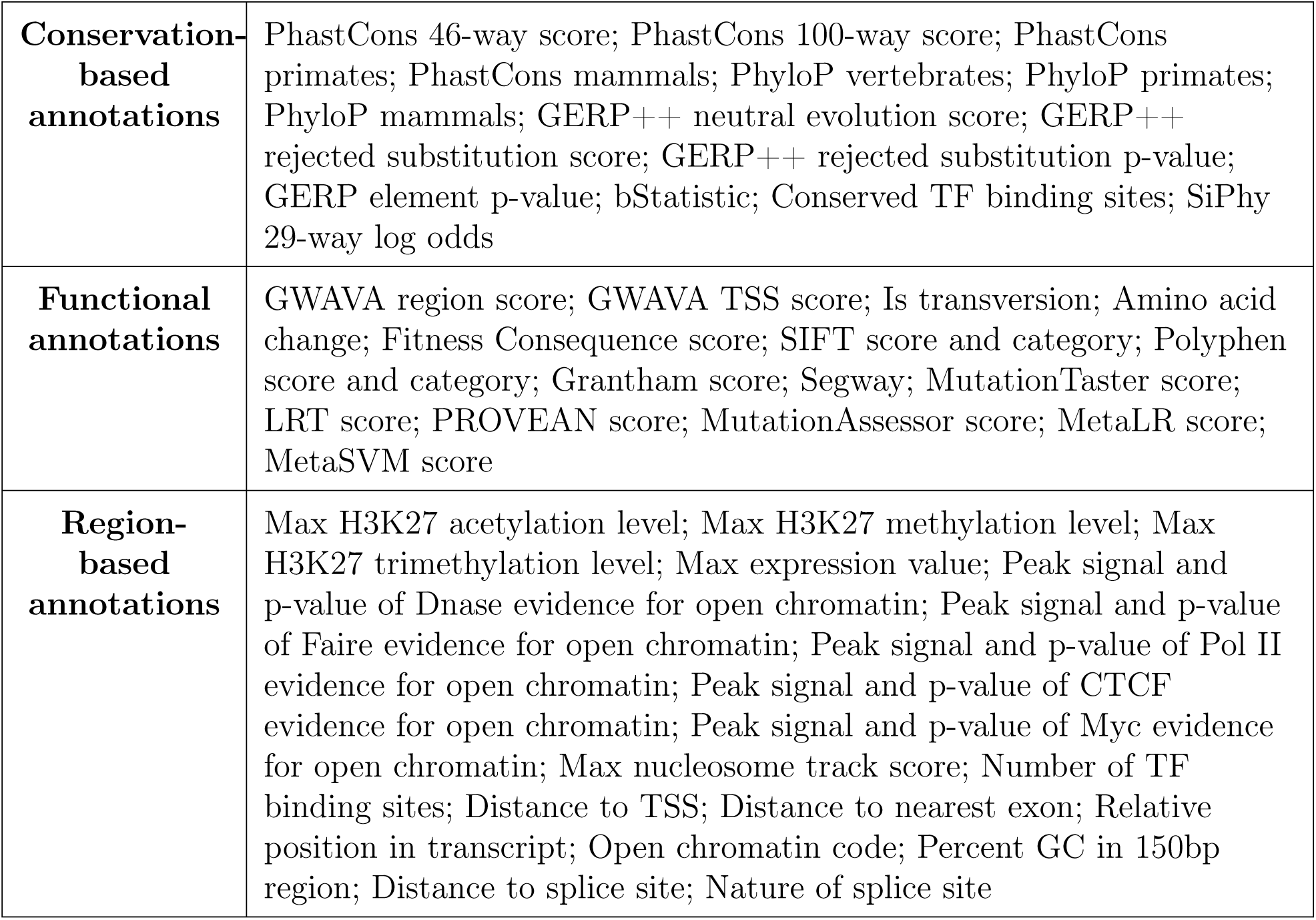
The list of annotations that were used as features within sORF-c.

## Discussion

Our ability to collect genomic data has now exceeded our ability to interpret this data. As whole genome sequencing becomes quick and cheap enough to be adopted for use in nonspecialist research laboratories worldwide, there is a need to develop tools that can provide useful insights from the unprecedented data that will be generated. Even while this work was being completed, the coordinates of a great many more potentially translated sORFs were made available on sORFS.org, following the completion of ribosome profiling experiments on various other human cell lines. This project demonstrates a workflow that could be implemented to quickly gain useful information on variants in sORF regions, or other non-coding regions of interest. It also highlights some of the limitations of currently available tools.

The performance of commonly used variant callers is highly dependent upon the genomic region, as demonstrated by the comparison presented here. Haplotype Caller calls variants most accurately across the whole genome, which agrees well with a previous systematic comparison of variant callers in exon-specific regions (Hwang *et al*, 2015). However, FreeBayes outperforms Haplotype Caller when deployed on sORFs. This may be because FreeBayes calls variants more accurately in the intergenic regions, which contain a higher density of problematic repeats. The large decrease in precision for each set of variants called in sORF regions is also partially due to the poor coverage of the truth VCF in repeat regions. Problematically, the GIAB VCF is the primary truth dataset used by the genomics community as a reference. This highlights the need for new consensus data with improved coverage in non-coding regions, which could be used to optimise new variant callers for these unexplored genomic regions.

By taking the intersection of variants identified by these variant callers, mutations that were not present in the truth VCF were likely identified, while false positives were minimised. Taking the intersection of variant calls represents a potentially useful step for future studies that require a high-precision list of variant calls, because variant calling algorithms cannot yet provide this within all non-coding regions. As regions of the human genome previously dismissed as non-coding continue to emerge as functionally important elements, these findings also demonstrate the importance of extending the coverage of high-performing variant calling algorithms to the non-coding regions.

Despite the high precision and sensitivity of the pathogenicity predicting algorithm demonstrated here, sORF-c requires significant improvements before it is useful to the genomics community. First and foremost, the classifier is likely overfitting the dataset, despite parameter optimisations to minimise this. sORF-c is highly accurate when predicting the pathogenicity of variants with the same distribution of metrics as the training data, but could likely perform less significantly on new clinical data obtained. Indeed, the demonstrated error rate of 11% on variants classified as ‘probably’ benign or pathogenic corroborates this; although this error rate may also have been affected by the misclassification of some variants with these classifications.

Obtaining a dataset of variants with sufficient depth to train a machine-learning algorithm proved challenging; the large list of benign variants taken from aligning the current reference genome to the inferred ancestral genome represented the best available dataset. Nonetheless, using this as the source of the majority of benign variants has likely biased the rules the classifier has learnt to differentiate pathogenic variants from benign. This ascertainment bias is most likely the cause of the reduced accuracy of sORF-c on variants obtained by more conventional means, such as manually curated databases like ClinVar. This hypothesis is supported by the substantially different distribution of feature scores observed between the two datasets of benign variants. Clinically evaluated databases of benign variants will instead be sought for future improvements to sORF-c.

Additionally, whilst the classifier presented here is termed a pathogenicity predictor, variants classified as pathogenic should not be construed as certainly causing the associated disease if present. Instead, the classifier should be interpreted as a variant prioritisation algorithm, whereby all identified variants are scored, and those with a higher score should be prioritised for investigation, as they are more likely to be causally linked to disease. An important next step for this project is therefore to calibrate the classifier with a larger number of known disease-causing variants, in order to identify precise thresholds in the scoring system that can more accurately describe the degree of prioritisation required. Nonetheless, by demonstrating that it is possible to train a classifier to detect sORF specific features, and that these features provide useful information for differentiating between pathogenic and benign variants, this project acts as a proof of concept for future progress in decoding the non-coding regions of the human genome.

## Materials and Methods

All databases and analyses performed used the GRCh37/hg19 assembly of the reference genome, which was completed in 2009.

### Data processing in the variant calling pipeline

Reads for the NA12878 genome were obtained from the Precision FDA website (https://precision.fda.gov/challenges/truth), and were produced by 50x paired-end sequencing on an Illumina HiSeq 2500 sequencer. The read length is approximately 148bp, and the insert size is approximately 550bp. The set of gold-standard variant calls, along with the BED file containing the coordinates of high-confidence regions, was downloaded from the Precision FDA website, but is also freely available elsewhere online (see (Zook *et al*, 2016)).

All variant calling tools were run on the Seven Bridges Genomics platform (https://www.sbgenomics.com/). Although some GATK tools were already present on this platform, the majority had to be created using a Docker image. The scripts for these were created locally, then uploaded on to the platform for use; they are now freely available to all platform users. The bash scripts that create Docker images of these tools can be found on GitHub account (www.github.com/fojackson8).

Raw sequence reads were first aligned to the reference genome (hg19) using the Burrows-Wheeler alignment algorithm (Li and Durbin, 2009). Because all popular read mapping algorithms operate on each read independently, it is impossible to globally minimise mismatches across all reads on the first pass. The indel realigner performs local realignment on small, potentially misaligned regions that occur as a result of an undetected indel.

The variant calling algorithms all assess the probability of a particular read signal being a true variant based on various base quality and context-specific metrics. However, these metrics are affected by artifacts from the sequencing experiment. For example, the PCR amplification step of the sequencing experiment results in allele amplification; if undetected, this can be mistakenly interpreted as multiple alleles with the same genotype, thus validating that genotype. It is therefore necessary to mark these duplicates, so they are not mistaken during subsequent variant calling. Variant calling al gorithms also rely on individual base quality scores, which are assigned during sequencing, and convey the likelihood of each call being correct. However, these quality scores co-vary significantly with sequencing method, and even between cycles of the same sequencing experiment, so must be recalibrated with the GATK Base Recalibration tool. This algorithm analyses the covariation between base quality scores and base context of known variants, and will recalibrate base quality scores based on this covariance.

The processing tools used were all taken from Picard tools or the Genome Analysis ToolKit (GATK), which were developed at the Broad Institute; tool documentation and guidelines can be found online at (https://software.broadinstitute.org/gatk/) and in the following paper (DePristo *et al*, 2011).

### Calling variants

Four variant calling algorithms were then tested on the same realigned, recalibrated BAM file. Haplotype caller and Unified Genotyper are both part of the Broad’s GATK, but use different read parsing and decision-making algorithms. Unified Genotyper is a naive Bayesian genotyper, which evaluates the posterior probability of the genotype at each locus based on the pileup of bases at that position and their associated quality scores. The likelihood, L, of a particular genotype, G, is calculated as follows:

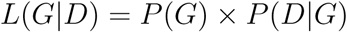

where G is the genotype corresponding to that variant and D is the data observed, which in this case is a set of base-specific quality metrics.

By contrast, Haplotype Caller performs local *de novo* assembly of haplotypes in variant-containing regions. This algorithm first identifies regions within aligned reads likely to contain variants, then reassembles these regions using a De Bruijn-like graph. Each haplotype is then realigned to the reference genome, and the likelihood of each genotype is calculated using Bayes’ rule as above. This local reassembly step improves Haplotype Caller’s accuracy of calling indels, and generally it is reported to be a more accurate variant caller. Both were included to verify this finding on this dataset. Both Unified Genotyper and Haplotype Caller are fairly aggressive callers, causing the resulting VCF to be enriched in false positives. This is because the best-practices pipeline involves a post-call variant quality score recalibration step, in which the majority of these false positives are removed using a Gaussian-mixture model trained on truth data such as HapMap, with pre-calculated quality metrics.

Platypus and FreeBayes do not require similar post-call variant recalibration. FreeBayes applies a similar Bayesian, haplotype approach to calling variants, but with adjustments to facilitate consideration of non-uniform copy numbers and multiallelism at each locus (Garrison and Marth, 2012). A single hard filter was applied to FreeBayes calls to reduce the number of false positive calls: all variants with a Phred-scaled quality score of less than five were filtered out. Platypus is a more recent variant calling algorithm designed to achieve similar performance to other variant callers with a less computationally demanding and therefore quicker method. Platypus does not map individual haplotypes to the reference genome, but instead performs reference-free sequence assembly, building de Bruijn graphs and searching them for polymorphisms (Rimmer *et al*, 2014).

The intersection of called variant sets was taken by first converting all variants into their primitive representation, using vcfallelicprimitives from the vcflib package. The intersection was then taken using vcfintersect, again from vcflib (https://github.com/vcflib/vcflib).

### Collecting and annotating known variants

Benign variants identified by aligning to the EPO 6-way primate alignment were downloaded with permission from Martin Kircher; for full details of how this dataset was constructed see Supplementary Material of (Kircher *et al*, 2014). The full HGMD database was kindly shared by Dr. David Cooper’s team at the Cardiff University; and the complete ClinVar database (clinvar_20170130) was downloaded from the publicly available ftp site. Figure 8 shows how the variants in each database were distributed across the genome. The subset of variants in sORF regions were found using tabix on bash, and each dataset was individually partitioned into training and test data (before combining), to ensure a similar proportion of pathogenic variants was present in both training and test sets.

**Figure 8:**
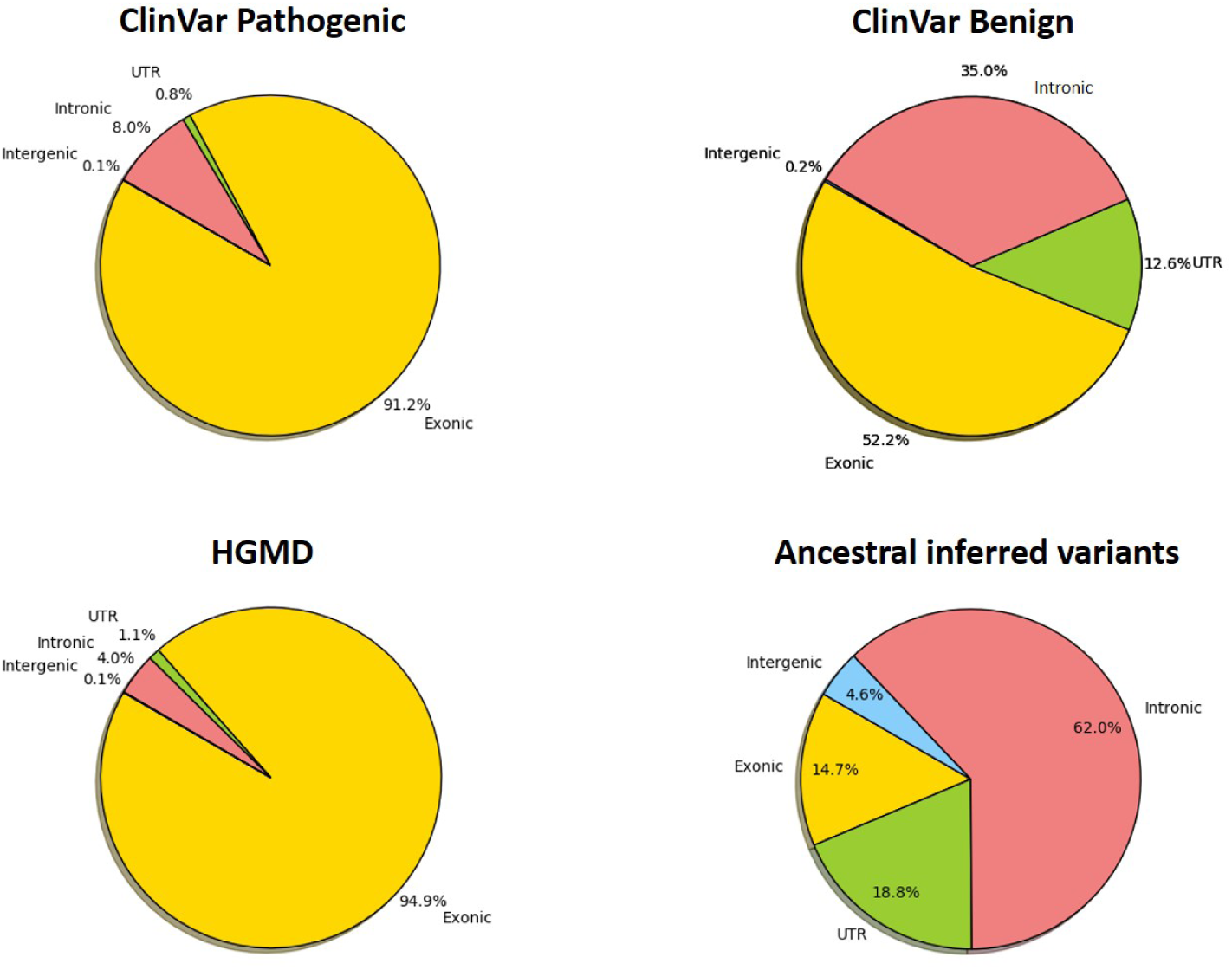
The distribution across the genome of variants from each dataset. Variants from the different benign and pathogenic databases differed significantly in their distribution across the genome. Each region is defined as in Figure 3.

All variants were then annotated using the annotate_variants.pl script within the Annovar tool (https://github.com/WGLab/doc-ANNOVAR/), which was installed on the Cambridge University Darwin Cluster. Annotations were downloaded using the Annovar download database command, and more information on each annotation used can be found at (http://annovar.openbioinformatics.org/en/latest/user-guide/download). Additional annotations were also taken from the CADD online scoring tool, which can be found at (http://cadd.gs.washington.edu/score). A complete list of annotations used is presented in Table 5.

### Handling missing values

Numerous variants in the training and test datasets had missing values for a subset of all feature scores. These missing values were all set to zero, because removing every variant that had a single missing annotation would have resulted in a severely depleted training set. Moreover, many of the annotations should only be present on a subset of the variants: for example, SIFT and Polyphen scorers both predict the effect of a variant on the resulting protein structure; the many variants outside of exonic regions should therefore have no score here. Assigning these missing values to zero represents an easily reproducible step that still allows useful information to be extracted at these positions.

### Parameter tuning for the Random Forests classifier

The RF model was implemented in Python 3, using the SciKit-learn (sklearn) package (http://scikit-learn.org/stable/). Although the RF classifier requires relatively little parameter tuning compared to other supervised machine-learning algorithms, parameters still need to be optimised to find the best compromise that maximises accuracy and minimises overfitting. A grid search over parameter space with cross-validation was performed using sklearn’s GridSearchCV. The grid search algorithm performed an exhaustive search within a stipulated range for each parameter listed in Table 6, using k-fold cross validation to evaluate performance at each step. The number of decision trees used in this forest was not optimised using a grid search; the value of this parameter largely depends on the desired trade-off between accuracy and computation time. Instead, the number of trees was increased until error rate stabilised (at which point further increases did not improve classification accuracy). 200 decision trees proved sufficient.

**Table 6:** The Random Forests tuning parameters that were optimised using Scikit learn’s GridSearchCV algorithm.

| Parameter | Range evaluated | Explanation | Optimal value |
| --- | --- | --- | --- |
| Max features | 2-15 | The maximum number of features to consider for each decision tree | 10 |
| Min samples per split | 2-10 | The minimum number of samples that each split of a node should possess | 2 |
| Min samples per leaf | 1-20 | The minimum number of samples required at each leaf. The smaller the number, the more prone each decision tree is to pick up noise. | 3 |
| Bootstrap | True or False | Whether to use bootstrap sampling of data when building decision trees | False |
| Evaluation criterion | Gini Impurity or Entropy | The function used to measure the quality of a split | Gini Impurity |
| Max depth | 3-None | The maximum depth of the tree. If none then each node will be expanded until all leaves contain less than min samples per split | None |

## Acknowledgements

Sudhakaran Prabakaran is a India-DBT Cambridge Lecturer. We thank the Seven Bridges Genomics team for making their cloud platform available to implement the variant calling pipeline. We thank all the members of Prabakaran Lab for comments on the manuscript. We thank Dr. David Cooper and Mr Matt Hayden from the Institute of Medical Genetics for sharing the HGMD database.

## URLs

Variant calling tools were run on Seven Bridges cloud-platform (https://www.sbgenomics.com/). vcfintersect tool was used from vcflib package (https://github.com/vcflib/vcflib). Benign training dataset was downloaded from (http://krishna.gs.washington.edu/members/mkircher/download/CADD/v1.3/training_data). Variant annotations were downloaded using the Annovar download database command from (http://annovar.openbioinformatics.org/en/latest/user-guide/download). Additional variant annotations were taken from the CADD online scoring tool (http://cadd.gs.washington.edu/score). SciKit-learn (sklearn) package was downloaded from (http://scikit-learn.org/stable/). All other scripts can be found at (www.github.com/fojackson8).

## Author contributions

SP designed the research. FJ and MW wrote the scripts. SP, FJ, and MW analyzed the data. SP and FJ wrote the manuscript with inputs from MW.

